# A Novel Method to Simultaneously Estimate Bacterial Respiration and Growth from Oxygen Dynamics

**DOI:** 10.64898/2025.12.10.693385

**Authors:** Ilgaz Cakin, Rebecca Millington, Samraat Pawar, Angus Buckling, Nicholas Smirnoff, Daniel Padfield, John Duffy, Gabriel Yvon-Durocher

**Author notes:** Corresponding author at: Living Systems Institute, University of Exeter, Stocker Road, Exeter, EX4 4QD, UK. Notes: The authors declare no competing financial interests.

## Abstract

Bacterial growth and respiration are fundamental metabolic processes that affect how energy is used and impact carbon sequestration at the ecosystem scale. However, these traits are usually quantified independently, growth is quantified with endpoint biomass measurements while respiration is quantified by monitoring oxygen or carbon dioxide. Because the two physiological traits are collected at different temporal and volumetric scales (hours-to-days for growth versus minutes-to-hours for respiration), reconciling them is challenging and often introduces scale-mismatch bias, obscuring causal links between metabolism and environmental drivers. In this study, we develop a novel method for quantifying the rates of bacterial growth and respiration concurrently from a single dissolved oxygen time series. Our approach introduces a model that couples exponential biomass growth with biomass-specific respiration, enabling the simultaneous inference of both rates in real time. We applied our high throughput method to 15 bacterial strains isolated from natural environments. Our approach yielded growth estimates in close agreement with measurements based on popular methods using optical density or flow cytometry (*R*^2^ > 0.9) with no evidence of taxon-specific bias. We also tested our approach in quantifying the effects of temperature on respiration, growth and carbon use-efficiency in *Pseudomonas* sp. Our method yielded typical unimodal thermal response curves for growth and respiration where rates were highest at moderate temperatures, while carbon-use efficiency increased from cooler temperatures, peaked around the thermal optimum (∼30–35 °C), and declined at high temperature. By quantifying respiration and growth simultaneously and in high throughput, our approach effectively enables measurement of microbial metabolic strategies and adaptations to stress. It offers a non-invasive and scalable tool for high throughput phenotyping and studies of environmental perturbations, enabling a new class of trait-based microbial ecology that links cellular physiology to broader ecosystem function.

## 1. Introduction

Microbial growth and respiration are essential physiological processes that drive the cycling of carbon and energy in ecosystems (Metze et al., 2023). By decomposing organic matter and fixing carbon into biomass, microbial communities play a key role in determining the distribution of carbon between biological pools and atmospheric CO₂ and, as such, in affecting biogeochemical fluxes at both local and global levels (Cao et al., 2021). Bacteria comprise a significant fraction of total ecosystem biomass (Fierer & Lennon, 2011) and are a major component of community-level respiration (McCallister et al., 2006), emphasising their critical contribution to ecosystem processes (Shi et al., 2019; Xu et al., 2018). At the cellular level, bacterial metabolic processes involve the allocation of energy between growth, leading to net biomass accumulation, and maintenance, which sustains necessary processes for cellular integrity (Pirt, 1965; Russell & Cook, 1995). This balance varies across environmental conditions, physiological states, and stressors, shaping microbial carbon use and energetic efficiency (Geider & Osborne, 1989; Shuter, 1979). Empirical observations indicate that total respiration comprises (i) a maintenance component that is independent of the instantaneous growth rate but varies with factors such as temperature, and (ii) a growth-associated component that is proportional to biomass synthesis (Geider & Osborne, 1989; Pirt, 1965). Understanding these components together is critical for linking cell metabolism to broader carbon fluxes at the ecosystem scale.

Despite this importance, growth and respiration are typically measured using distinct, asynchronous methods; oxygen uptake or CO₂ release for respiration, and optical density, cell counts, or biomass yield for growth (Warkentin et al., 2007). When research questions aim to link growth and metabolism, for example by attempting to understand how environmental variation affects energy allocation, measuring growth and respiration across different scales can make reconciling these rate measurements challenging. For example, rates of growth measured via cell counts or biomass change over time are typically measured over timescales of several hours to days, while rates of respiration are measured over minutes to hours. Chemostats can circumvent some of these challenges by measuring growth and metabolic flux *in situ* at steady state, however chemostats are limited in their scale both in terms of experimental replication (numbers of treatments x replicates) and timescale (long-term contamination free chemostats are challenging to maintain) (Miller et al., 2013; Plouchart et al., 2018). These factors limit our understanding of the dynamic interplay between respiration and growth rate, particularly under changing environmental conditions. Recent work showed that asynchronous measurement of respiration and biomass production can distort carbon budgets, systematically overestimating respiration, underestimating growth efficiency, and inflating the apparent role of bacteria in community metabolism (Guo et al., 2022). Environmental factors such as temperature, pH, nutrient availability, and osmotic stress influence growth and respiration through distinct physiological mechanisms. As a result, integrated, time-resolved measurements that capture both processes are increasingly essential for understanding microbial energy partitioning. Decoupling between these processes at the cellular level can scale up to affect ecosystem functions such as carbon sequestration, nutrient turnover, and greenhouse gas fluxes (Knapp & Huang, 2022; Russell & Cook, 1995).

Here we present a novel method to infer bacterial growth rates and respiration simultaneously from a single time series of oxygen concentration measurements. We developed a mathematical model that treats growth as an exponential process and assumes that oxygen uptake is proportional to biomass. We applied our approach to estimate growth and respiration in 15 phylogenetically diverse bacterial strains under standardised laboratory conditions. Growth rates derived from our novel oxygen gradient method were confirmed by independent measurements of growth rate made using flow cytometry and spectrophotometry. To illustrate its broader utility, we applied our approach to *Pseudomonas* sp. exposed to a controlled thermal gradient, enabling the measurement of temperature-dependent changes in growth, respiration, and carbon use efficiency from a single oxygen time series. This unified approach reveals new insights into microbial energy allocation and metabolic flexibility across conditions, offering valuable tools for both basic ecology and applied microbiology.

## 2. Materials and Methods

### 2.1. Bacterial taxa

Experiments were conducted using 15 heterotrophic bacterial taxa originally isolated from Icelandic geothermal pools (García et al., 2018). Taxa were identified by 16S rRNA gene sequencing and included *Aeromonas* sp. (strain A and B), *Bacillus* sp., *Pseudomonas* sp., *Flavobacterium* spp. (strain A and B), *Yersinia* sp., *Serratia* sp., *Burkholderia* sp., *Buttiauxella* sp., *Chryseobacterium* sp., *Herbaspirillum* sp., *Arthrobacter* sp., *Chromobacterium* sp., and *Acinetobacter* sp.

### 2.2. Bacterial monoculture preparations

Each bacterial isolate was revived from −80 °C glycerol freezer stocks and streaked onto LB agar plates. A single colony was then inoculated into sterile LB medium (Lysogeny Broth; 10 g tryptone, 5 g yeast extract, 10 g NaCl per litre) and incubated at 30 °C for 16 hours (Panasonic Corporation, Japan). Following incubation, optical density at 600 nm (OD₆₀₀) was measured by Infinite 200 PRO plate reader (Tecan Life Sciences, Switzerland), and cultures were diluted into fresh, pre-warmed, and aerated LB medium to an initial OD₆₀₀ of 0.05. Following a 2-hour incubation to promote entry into exponential growth, cultures were further diluted to a final OD₆₀₀ of 0.0005.

### 2.3. Dissolved oxygen measurements

Oxygen consumption was measured using SensorDish^®^ readers (PreSens GmbH, Germany), which enable real-time, non-invasive monitoring of up to 24 samples per plate via optical sensor spots containing a fluorescent dye quenched by oxygen (Werner et al., 2021). The degree of quenching correlates with the oxygen partial pressure, enabling non-invasive, real-time measurement of dissolved oxygen concentrations (PreSens Precision Sensing GmbH., 2025). The system was factory-calibrated prior to use, and before each experimental run we additionally performed a two-point calibration against 0% and air-saturated oxygen following the manufacturer’s protocol, to verify linearity and stability of the sensor response over the range relevant for our assays. Vials were sterile upon first opening, as supplied by the manufacturer, and were handled aseptically throughout. All liquid handling was performed in a sterile laminar flow hood to prevent contamination. Five replicate vials were prepared per strain by aliquoting from the same pre-grown culture at OD₆₀₀ = 0.0005. Each 5 mL vial was filled to slight convexity, just above the rim, to eliminate headspace and thereby minimise atmospheric *O*_2_ exchange, then sealed with the manufacturer-supplied gas-tight caps. Vials were placed on the readers in temperature-controlled incubators set to a constant temperature (30 °C for the multi-taxon experiment and fixed set points between 20 and 40 °C for the temperature experiment) and incubated for approximately 45 minutes to allow the SensorDish^®^ optical signal and baseline to stabilise; during this period the reported dissolved-oxygen values may drift slightly upward as the reader optics warm up and internal temperature-compensation and calibration routines settle. Dissolved oxygen concentrations were then recorded at 1-minute intervals over a ∼200-minute period.

### 2.4. Optical density and flow cytometry measurements

To obtain initial cell densities without disturbing the oxygen measurements, we prepared a parallel set of cultures using the same inocula, vials, and equipment, and incubated them under identical environmental conditions in a separate incubator. This incubator was the same brand and model as the one housing the SensorDish^®^ unit and was set to the same temperature; temperatures in both incubators were monitored regularly and showed no systematic differences. For each taxon, five parallel vials were prepared. After 45 minutes of incubation (matching the SensorDish^®^ stabilisation period), subsamples were taken from all five vials, fixed immediately, and later quantified by flow cytometry. These parallel cultures provided the taxon-specific initial (*t*_0_) cell densities used together with OD₆₀₀ and flow-cytometry measurements.

In the SensorDish^®^ experiment itself, subsamples were taken from each vial at the time of inoculation and quantified by flow cytometry to determine the inoculated cell density (*N*_inoc_) for each taxon × replicate. These *N*_inoc_values were later used to reconstruct replicate-specific *N*_0_ at the start of each dissolved-oxygen time series. Final OD₆₀₀ and flow-cytometry measurements at *t*₁ were obtained directly from the corresponding vials within the SensorDish^®^ incubator, which remained sealed during the run to maintain uninterrupted oxygen monitoring. Thus, *t*_0_cell densities were estimated from the parallel cultures, whereas *N*_inoc_ and *t*_1_cell densities were measured from the same SensorDish^®^ vials that generated the oxygen time series.

Optical density at 600 nm (OD₆₀₀) was measured using a microplate reader to estimate total biomass. For flow cytometry, 490 µL of culture was mixed with 10 µL of 25% glutaraldehyde (Sigma-Aldrich, USA), incubated at 4 °C for 20 minutes, and then snap-frozen in liquid nitrogen. Samples were stored at −70 °C until further analysis. Prior to flow cytometric measurement, samples were thawed at 37 °C with shaking at 300 rpm for 5 minutes, then diluted by mixing 50 µL of thawed culture with 450 µL of sterile phosphate-buffered saline (PBS). SYBR Green I (Thermo Fisher Scientific, USA) was used as a nucleic acid stain and added by mixing 2.5 µL of a 100X solution into each 500 µL sample. Samples were incubated in the dark at room temperature for 1 hour prior to analysis. A CytoFLEX S flow cytometer (Beckman Coulter, Inc., USA) was used to quantify cell counts. SYBR-stained sterile LB was used as a blank to determine background fluorescence.

### 2.5. Oxygen model fitting for respiration and growth rate estimates

To estimate microbial growth and respiration simultaneously from dissolved-oxygen time series, we developed a model that links exponential increases in biomass to cumulative oxygen consumption. We assume exponential growth of cell numbers, so that

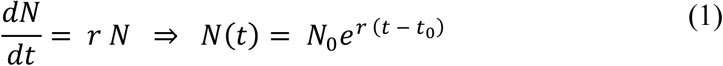

where *N*(*t*) is cell density (cells L⁻¹) at time *t*, *N*_0_ is the density at time *t*_0_, and *r* is the specific growth rate (min⁻¹).

Assuming a constant per-cell respiration rate *R* (mg O₂ cell⁻¹ min⁻¹), the instantaneous rate of oxygen consumption is proportional to biomass,

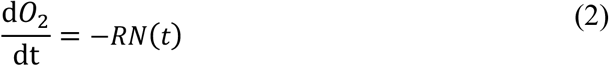

where *O*_2_ is the dissolved oxygen concentration (mg L⁻¹). Combining (1) and (2) and integrating from *t*_0_to *t* gives

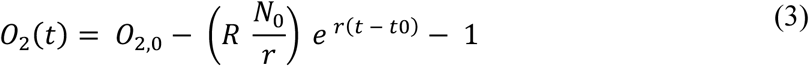

where *O*_2,0_= *O*_2_(*t*_0_) is the oxygen concentration at the start of the fitted trajectory. In this expression, *O*_2,0_ sets the baseline, the factor (*RN*_0_/ *r*) controls the overall amplitude of the decline, and *r* controls its curvature.

Prior to fitting, raw oxygen time series were pre-processed as follows. First, we excluded the initial stabilisation period of the SensorDish^®^ measurement (approx. 45 min). Second, each series was automatically trimmed to retain only the portion of the trajectory from the post-stabilisation peak (the local maximum or “tip” where oxygen first begins to decline) to the point at which the decline in oxygen concentration starts to decelerate (i.e. where the absolute rate of decrease, |dO_2_/dt|, begins to fall), removing the late, near-stationary part of the trajectory where exponential growth is no longer a reasonable approximation. Time was then re-zeroed so that the first retained point corresponded to *t* = 0. Finally, we normalised each series by its initial oxygen level in the trimmed window, so that

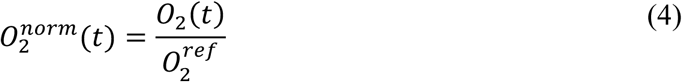

where *O*_2_*^ref^* is the mean of the first three oxygen measurements in the trimmed series. This pre-processing ensures that the model fit is driven primarily by the shape of the oxygen decline rather than by small errors in absolute calibration, and removes any apparent initial increase in dissolved oxygen caused by instrument drift so that it cannot influence the estimated parameters.

With *t*_0_ set to 0, equation (3) can be written in terms of the normalised oxygen as

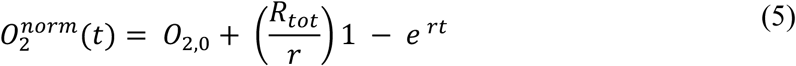

Here *O*_2_*^norm^*^/^(*t*) is the normalised dissolved oxygen, *O*_2,0_is a dimensionless offset that allows the initial normalised oxygen to deviate slightly from 1, and *R_tot_* is a total respiration scaling parameter that, in the underlying dimensional form, corresponds to the product *RN*_0_. In the fitting step, *N*_0_ does not appear explicitly: we set *N*_0_= 1 in the model, so *RN*_0_is absorbed into *R*_tot_. Replicate-specific *N*_0_values and per-cell respiration rates are recovered later using inoculation densities and the fitted growth rate.

The model in equation (5) was implemented as a custom R function and fitted separately to each taxon x replicate time series by nonlinear least squares using the nlsLM function from the minpack.lm package. For each series we estimated *r* (per min), *R_tot_*, and *O*_2,0_, with time *t* expressed in minutes from the start of the trimmed trajectory. Starting values were taken from the data (an initial estimate of *r* from the early-phase slope of log-transformed, normalised oxygen, and the initial rate of decline for *R_tot_*). During fitting, parameters were constrained to broad, physiologically reasonable ranges to avoid clearly non-physical solutions: specifically, 10^-4^ <= *r* <= 10^-1^ per min, 10^-5^ <= *R_tot_* <= 5.10^-1^, and 0.8 <= *O*_2,0_<= 1.2. These intervals act as physiological guardrails that prevent the optimiser from wandering into unrealistic regions of parameter space, rather than forcing the fits towards particular values. After an initial fit, points with residuals larger than two standard deviations for that series were treated as outliers, removed, and the model was refitted. Goodness of fit was summarised using a pseudo-*R*^2^statistic, calculated as

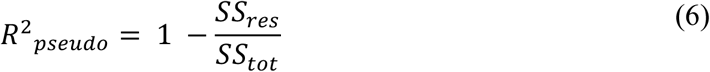

where *SS_res_* is the residual sum of squares from the fitted model and *SS_tot_* is the total sum of squares of the normalised oxygen values around their mean. Fits with low *R*^2^*_pseudo_*, large relative standard errors, or non-physiological parameter estimates were flagged and excluded from downstream analyses. To assess model adequacy, we visually inspected residuals for each taxon and replicate and did not observe any obvious temporal structure or strong heteroscedasticity; representative residual plots are provided in Supplementary **Fig. S1**.

To obtain replicate-specific initial cell densities at the start of the oxygen trajectories, we combined the measured inoculation densities with the *O*_2_-based growth rate estimates. Let *N_inoc_* be the cell density at inoculation (cells per µL), measured by flow cytometry from the SensorDish vials (Section 2.4), and *r* is the specific growth rate (per min) from the oxygen model. If there is a delay Δ*t* (min) between inoculation and the first oxygen measurement (Δ*t* = 45 min in our experiments), exponential growth implies

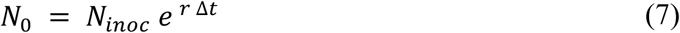

where *N*_0_ is the cell density (cells per µL) at the start of the oxygen time series (note: we multiply *N*_0_ by the factor 1e6 to convert from µL to L for downstream unit compatibility). This *N*_0_was then used to convert the total respiration scaling parameter from the oxygen model into a per-cell respiration rate. Specifically, if *R_tot_* is the total respiration scaling parameter (units min⁻¹) from equation (5), and *O*_2_*^ref^* is the initial dissolved oxygen concentration (mg *O*_2_ L⁻¹) used for normalisation in equation (4), the corresponding per-cell respiration rate *R* (mg *O*_2_ cell⁻¹ min⁻¹) is

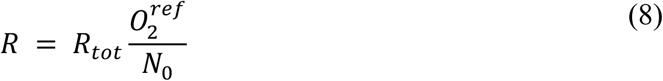

Thus, the oxygen-based model directly provides *r* and *R_tot_* for each replicate, and *N*_0_ and *R* are reconstructed from the inoculation densities and the known delay between inoculation and oxygen measurements.

### 2.6. Sensitivity and model validation analyses

In the oxygen model given in equation (5), the decline in normalised dissolved oxygen is parameterised in terms of the total respiration scaling parameter *R_tot_*, which does not include the initial biomass explicitly. The specific growth rate *r* is therefore determined directly from the curvature of the normalised oxygen decline and does not depend on the assumed value of *N*_0_, whereas the per-cell respiration rate *R* is obtained by combining *R_tot_* with the initial oxygen concentration and the initial cell density via equation (8). To quantify how uncertainty in *N*_0_ propagates into the per-cell respiration rate, we performed a Monte Carlo sensitivity analysis (Supplementary **Table S1**).

Using the inoculation-based estimates of *N*_0_ at the start of the oxygen measurements (equation (7)), we first obtained taxon-level summaries of *N*_0_(mean and standard deviation in cells L⁻¹). For each taxon × replicate oxygen time series, we then drew 300 realisations *N*_0_^k^ from a normal distribution with the corresponding taxon-specific mean and standard deviation, truncated so that*N*_0_^k^> 0. For each draw, we held the fitted values of *R_tot_* and *O*_0_*^ref^* fixed, and recalculated the per-cell respiration rate as

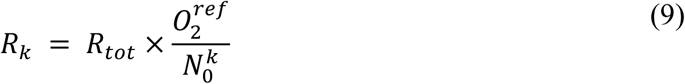

For each replicate, we summarised the resulting distribution of *R_k_* by its relative standard deviation (standard deviation divided by the mean), providing a direct measure of how strongly uncertainty in *N*_0_ influences the inferred per-cell respiration rates.

In addition, we carried out a synthetic-data validation to test whether both *r* and *R* can be reliably recovered from single dissolved-oxygen time series under realistic experimental conditions (Supplementary **Fig. S2**, **Table S2**). We generated artificial *O*_2_ trajectories from the oxygen model in equation (5), using parameter combinations spanning the empirical range of *N*_0_, *r*, and *R* observed in our experiments. These trajectories were sampled at realistic intervals (down to 1 min) over time windows up to approximately 3h, and Gaussian noise was added at levels comparable to or higher than the SensorDish^®^ system. Each synthetic dataset was then processed with exactly the same trimming, normalisation, and nonlinear fitting pipeline as the empirical data, and the recovered parameter estimates were compared directly with their known true values.

A visual summary of the experimental pipeline and analytical workflow is provided in **Fig. 1**. For additional practical guidance on implementing the oxygen-based model, including common pitfalls and recommended solutions, see Supplementary **Note S1**.

**Figure 1.**
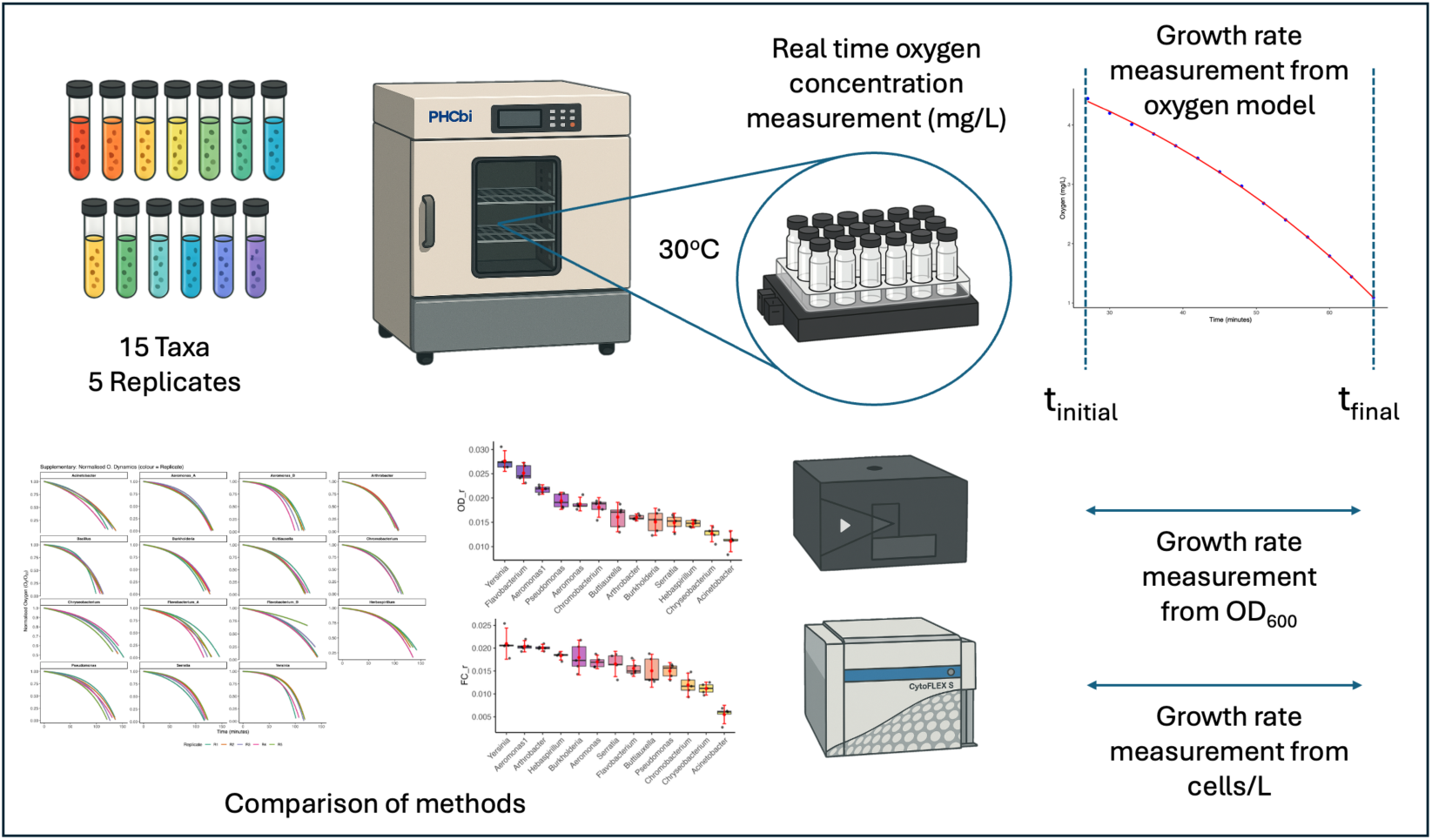
Overview of experimental design and methodology. Fifteen bacterial taxa were incubated in LB medium at 30 °C in vials monitored using SensorDish^®^ Readers for real-time oxygen concentration measurements. Growth rates were estimated from oxygen gradient model fits, optical density measurements, and flow cytometry. Comparative analyses across methods and taxa were conducted to evaluate method consistency.

### 2.7. Environmental gradient assays and stress response modelling

To investigate metabolic responses under environmental stress, we applied the oxygen-based modelling framework to a single strain of *Pseudomonas* sp. exposed to a controlled temperature gradient. Temperature effects were assessed by incubating cultures at 20, 24, 28, 32, 36, and 40 °C, with four replicates per condition. Cultures were incubated in sealed vials and monitored at 1-minute intervals for approximately 3h, so that rapid oxygen dynamics at temperatures deviating from the thermal optimum were captured. The cell density at inoculation (*N_inoc_*, cells µL⁻¹) was measured by flow cytometry. The specific growth rate *r* (min⁻¹) was then estimated from the dissolved-oxygen time series using the nonlinear oxygen model (equation 5). Given the known delay Δ*t* between inoculation and the first oxygen measurement, the initial cell density at the start of the oxygen trajectory (*N*_0_, cells L⁻¹) was reconstructed by back-extrapolating under exponential growth, as described in Section 2.5.

For the estimation of cellular carbon content, cell dimensions were based on *Pseudomonas* sp., described as rod-shaped bacteria measuring 0.5–0.8 µm in diameter and 1.5–3.0 µm in length (Moore & Flaws, 2011). The resulting mean cell volume was multiplied by 100 fg C µm^−3^, which lies at the upper end of values reported for actively growing bacterial cells (Braun et al., 2016). Specific growth rates (*r*, in min⁻¹) were then converted to carbon-based biomass production rates by multiplying *r* by the initial cell density (cells L⁻¹) and the estimated cellular carbon content (fg C cell⁻¹), yielding growth in fg C L⁻¹ min⁻¹.

Respiration rates were estimated as per-cell *O*_2_ consumption rates (*R*, in mg *O*_2_ cell⁻¹ min⁻¹) from oxygen-consumption curves using the nonlinear model (eq. 5). To express respiration rate in carbon units, first *R* was multiplied by the initial cell density (*N*_0_, cells L⁻¹), yielding volumetric oxygen consumption rates (mg *O*_2_L⁻¹ min⁻¹). These were converted to mol *O*_2_L⁻¹ min⁻¹ using the molecular weight of *O*_2_ (31.998 g mol⁻¹), then to fg C L⁻¹ min⁻¹ assuming a standard first-order approximation of a 1:1 molar ratio between *O*_2_ consumption and carbon oxidised for aerobic respiration of moderately oxidised substrates, and using the atomic mass of carbon (12 g mol⁻¹). These stoichiometric and carbon-density assumptions are used only to place growth and respiration on a common carbon scale in the *Pseudomonas* sp. temperature experiment.

Carbon-use efficiency (CUE) was calculated as the ratio of carbon-based biomass production (growth) rate, *P*, to the sum of biomass production and respiration rate, *P* + *R*

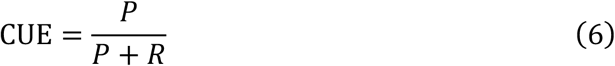

### 2.8. Data handling & statistical analyses

All statistical analyses were performed in R (version 4.4.3) (R Core Team, 2010). All analysis scripts are available at (https://github.com/icakin/OxygenModel). Replicate-level growth rates were first visualised using boxplots grouped by taxon and by method, created with *ggplot2* (Wickham & Sievert, 2009) and *patchwork* (Pedersen, 2019). This allowed assessment of both within-taxon variability and global differences across oxygen, OD₆₀₀, and FC-derived estimates. Growth rates from optical density and flow cytometry were estimated by calculating the exponential rate of increase between the initial and final timepoints of each replicate:

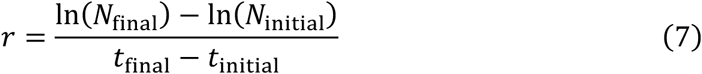

To evaluate agreement between oxygen-derived growth rates and those estimated from optical density (OD₆₀₀) and flow cytometry (FC), linear mixed-effects models were then applied using *lme4::lmer()* (Bates et al., 2015).

Bacterial taxon was included as a random intercept in the mixed-effects models to account for baseline differences in growth rates among taxa. This allowed us to test whether a consistent relationship was present across measurement methods while controlling for taxon-specific variation. Fixed-effect slopes were used to quantify the expected change in oxygen-derived growth per unit change in growth rates estimated from either OD₆₀₀ or flow cytometry (FC). Model performance was evaluated using marginal *R*^2^values, calculated via the *performance* package (Lüdecke et al., 2019), which indicate the proportion of variance explained by the fixed predictors. For visualisation, replicate-level data were normalised by subtracting taxon-specific random intercepts, using functions from *dplyr* (Wickham, 2015) and *tidyr* (Wickham et al., 2023). This procedure removed baseline offsets and enabled a clearer comparison of method agreement on a common scale.

To further assess systematic bias and agreement between methods, we applied Bland–Altman analysis (Bland & Altman, 1986). Bland–Altman plots are widely used to compare two quantitative measurement techniques because they visualise not only the average agreement but also potential biases across the measurement range. For each replicate, the difference between paired measurements was calculated as:

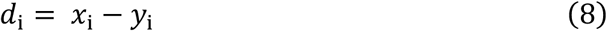

where *x*_i_ is the growth rate estimated from oxygen dynamics and *y*_i_ is the corresponding estimate from OD₆₀₀ or FC. The average of the two methods was computed as:

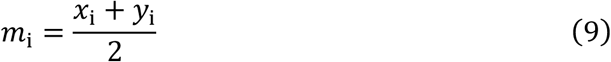

The Bland–Altman plot displays *d*_i_ on the y-axis against *m*_i_on the x-axis. The mean difference *d*^-^ provides an estimate of systematic bias between methods. Limits of agreement (LoA) were then calculated as:

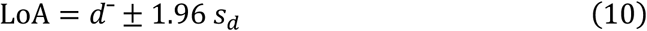

where *s*_d_ is the standard deviation of the differences. These limits represent the interval within which 95% of the pairwise differences are expected to fall, thereby quantifying the range of divergence between methods across the observed growth rate spectrum. All analyses were conducted at replicate-level resolution in order to preserve within-taxon variability rather than averaging across replicates.

Growth and respiration rates for the temperature-gradient experiments were estimated by fitting a nonlinear oxygen-consumption model to each replicate using *nlsLM()* from the *minpack.lm* package (Elzhov et al., 2016). Thermal responses of growth and respiration were then characterised by fitting Sharpe–Schoolfield models to the estimated rates, allowing identification of activation energies and thermal optima (Garcia et al., 2022; Schoolfield et al., 1981). Carbon use efficiency (CUE) was modelled with an Arrhenius function, reflecting its generally monotonic decline with increasing temperature (Padfield et al., 2016).

## 3. Results

### 3.1. Deriving bacterial respiration and growth rate from oxygen time-series

The model (eq. 5) effectively captured the temporal dynamics of oxygen consumption, enabling simultaneous estimation of specific growth rate (*r*) and the per-cell respiration rate (*R*) from dissolved oxygen time-series data (Fig. 2). The fitted curves closely followed the observed oxygen depletion trajectories across replicates and taxa, with a median pseudo-*R*^2^ **>** 0.99 (see Methods), indicating that the model reproduces the observed trajectories very closely. We use pseudo-*R*^2^ here solely as a summary of curve-fitting quality; potential deviations from exponential growth or stationarity are assessed separately via our trimming procedure, sensitivity analyses, and cross-method comparisons. Residuals showed no obvious temporal structure or strong heteroscedasticity (Supplementary **Fig. S1**), consistent with an adequate statistical description of the oxygen decline. Estimated growth rates ranged from 0.008 to 0.046 min⁻¹, while per cell respiration rates spanned from 1.53e-11 to 1.43e-9 mg O₂ cell⁻¹ min⁻¹, reflecting substantial inter-taxon differences under standard conditions. Full oxygen depletion curves for all five replicates of the 15 bacterial taxa are provided in Supplementary **Fig. S3.**

**Figure 2.**
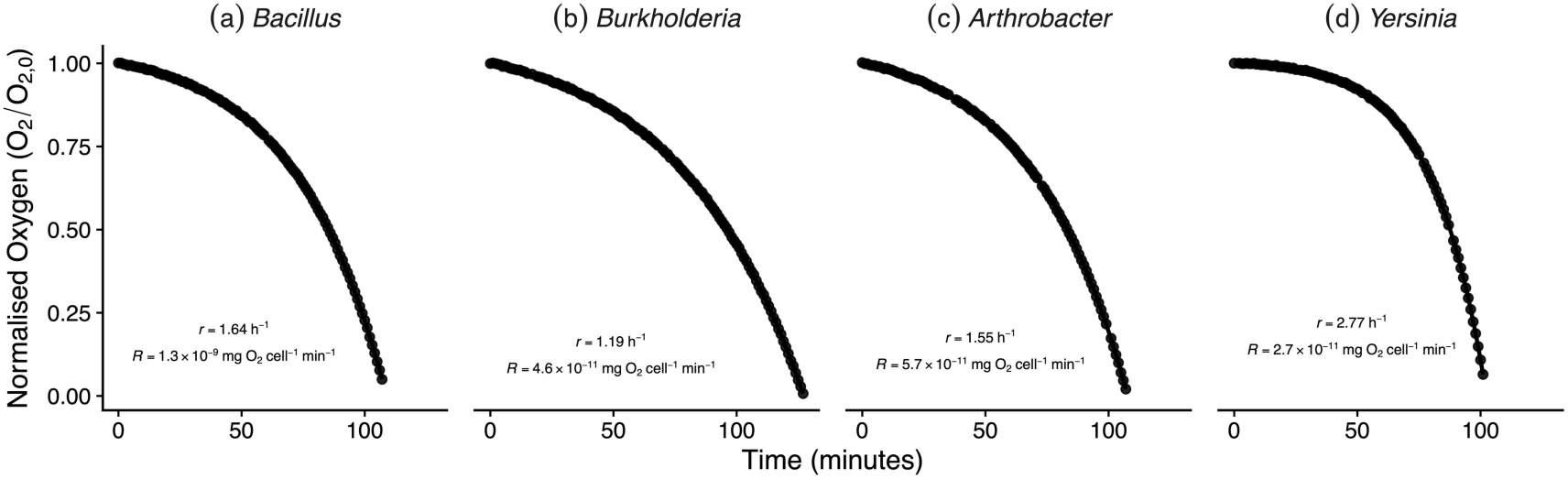
Taxon-specific oxygen consumption dynamics and model-derived metabolic rates. Dissolved oxygen concentration (mg L⁻¹) over time for four representative bacterial taxa (selected from 15 total) under standardised assay conditions. Black dots show observed oxygen concentrations; solid lines represent model fits. The mechanistic model simultaneously estimated specific growth rates per minute (*r*) and per-cell respiration rates per minute (*R*), reported within each panel.

### 3.2 Validating the modelling pipeline

To quantify how uncertainty in *N*_0_propagates into the estimation of the per-cell respiration rate, *R*, we performed a Monte Carlo sensitivity analysis. We found that the inferred *R* is moderately sensitive to uncertainty in *N*_0_. For most taxa, the median relative standard deviation of *R* was on the order of 0.1–0.3 (i.e. ∼10–30%), with larger values occurring only in a small subset of taxa where *R* was less tightly constrained by the oxygen time series (Supplementary **Table S1**). Note, the specific growth rate *r* is determined from the curvature of the normalised oxygen decline and therefore does not depend on the assumed value of *N*_0_.

Next, we carried out an analysis based on synthetic-data to validate our modelling procedure and test whether both *r* and *R* can be reliably recovered from single dissolved-oxygen time series using our modelling framework. We generated synthetic dissolved-oxygen time series from equation (5) for a grid of growth rates (*r*) and per-cell respiration rates (*R*) spanning the range observed in our experiments, using a fixed initial biomass *N*_0_. For each parameter combination, we added Gaussian observation noise in normalised *O*_2_ and varied the sampling interval and total duration to mimic our experimental designs. We found that our model-fitting pipeline recovered both parameters with negligible bias (relative bias generally <2%) and high accuracy (RMSE for *r* on the order of 10,^>^min^-1^ and mean pseudo-*R*^2^> 0.99 across all conditions, see Supplementary **Table. S2**). Parameter uncertainty increased systematically as we increased higher observation noise and, to a lesser extent, when we reduced the length of the time series (60 min) (Supplementary **Table. S2**).

Finally, we asked whether our parameter estimates might be biased by potential oxygen limitation at low dissolved *O*_2_. In our standard workflow, each time series is trimmed to the main depletion phase, which can extend into relatively low oxygen levels. To test the robustness of our results to this choice, we re-fitted the model for all strain–replicate combinations using only data points with normalised oxygen ≥ 0.5, thereby restricting the fit to the high-oxygen portion of the trajectories. The resulting growth and respiration estimates (*r*_0.5_, *R*_0.5_) were virtually indistinguishable from those obtained with our standard trimmed window (median *r*_0.5_/*r* = 0.9997, interquartile range 0.9974–1.0024; median *R*_0.5_/*R* = 1.0004, interquartile range 0.9964–1.0034; maximum absolute deviations ∼1.0% for *r* and ∼1.2% for *R*). This indicates that potential oxygen limitation at lower dissolved *O*_2_ does not materially bias our parameter estimates (Supplementary Table S3).

### 3.2. Comparison of growth rate estimates across measurement methods

Growth rate estimates obtained from the oxygen-based model were highly consistent with those derived from conventional approaches, including optical density (OD₆₀₀) and flow cytometry (FC), across all 15 bacterial taxa tested (Fig. 3). Oxygen-derived estimates spanned a very similar range to OD₆₀₀- and FC-based measurements, and distributions across taxa showed comparable medians and variability.

**Figure 3.**
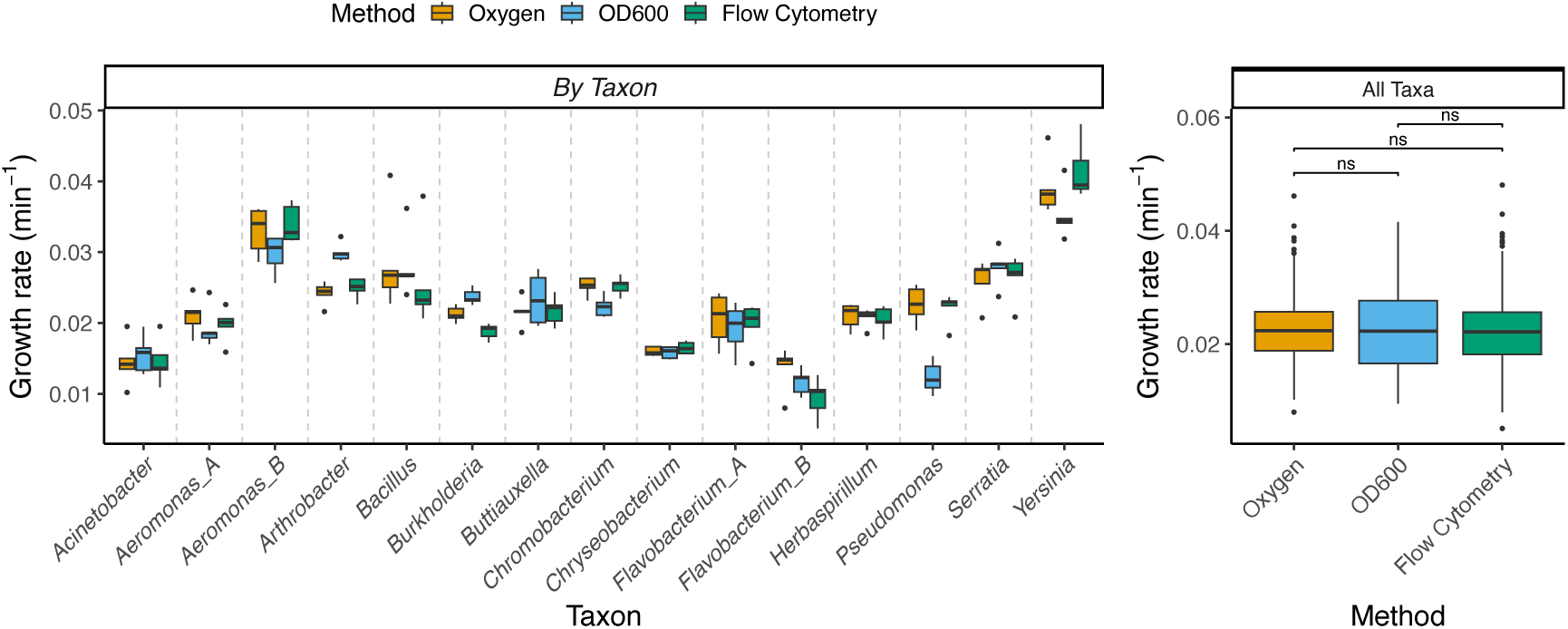
Comparison of bacterial growth rate estimates across methods. Growth rates (per minute) were estimated using three independent methods: oxygen consumption modelling (orange), optical density at 600 nm (OD₆₀₀; blue), and flow cytometry (green). The left panel shows method-specific growth rate distributions for each of the 15 bacterial taxa. Each boxplot represents replicate-level estimates, with horizontal lines indicating the median and whiskers denoting the interquartile range.

Some taxon-specific deviations were evident, with oxygen-based estimates occasionally higher or lower than those from OD₆₀₀ or FC, but these differences varied in direction across taxa and did not indicate systematic bias. When estimates were pooled across taxa, the three methods yielded nearly identical distributions. Visual patterns were supported by statistical analysis. A mixed-effects ANOVA with method as a fixed effect and taxon as a random intercept detected no significant overall effect of method, as a likelihood ratio test comparing models with and without method terms confirmed (*x*^2^= 2.60, df = 2, *p* = 0.27). Pairwise contrasts with Bonferroni correction likewise indicated no significant differences among oxygen-, OD₆₀₀-, and flow cytometry–derived growth rates (all ns).

To quantitatively compare growth rate estimates derived from oxygen consumption with those obtained via OD₆₀₀ and flow cytometry (FC), we fitted linear mixed-effects models incorporating taxon as a random intercept. Oxygen-derived growth rate was used as the response variable, and either OD₆₀₀- or FC-derived growth rate was used as the predictor. Oxygen-based estimates showed a strong linear relationship with OD₆₀₀-based growth rates (*R*^2^= 0.943), with a fixed-effect slope of 0.913 (95% CI: 0.764–1.062) and an intercept of 0.003 (Fig. 4A). The relationship between oxygen- and FC-derived growth rates was similarly strong (*R*^2^ = 0.968), with a slope of 0.887 (95% CI: 0.817–0.957) and an intercept of 0.003 (Fig. 4B).

**Figure 4.**
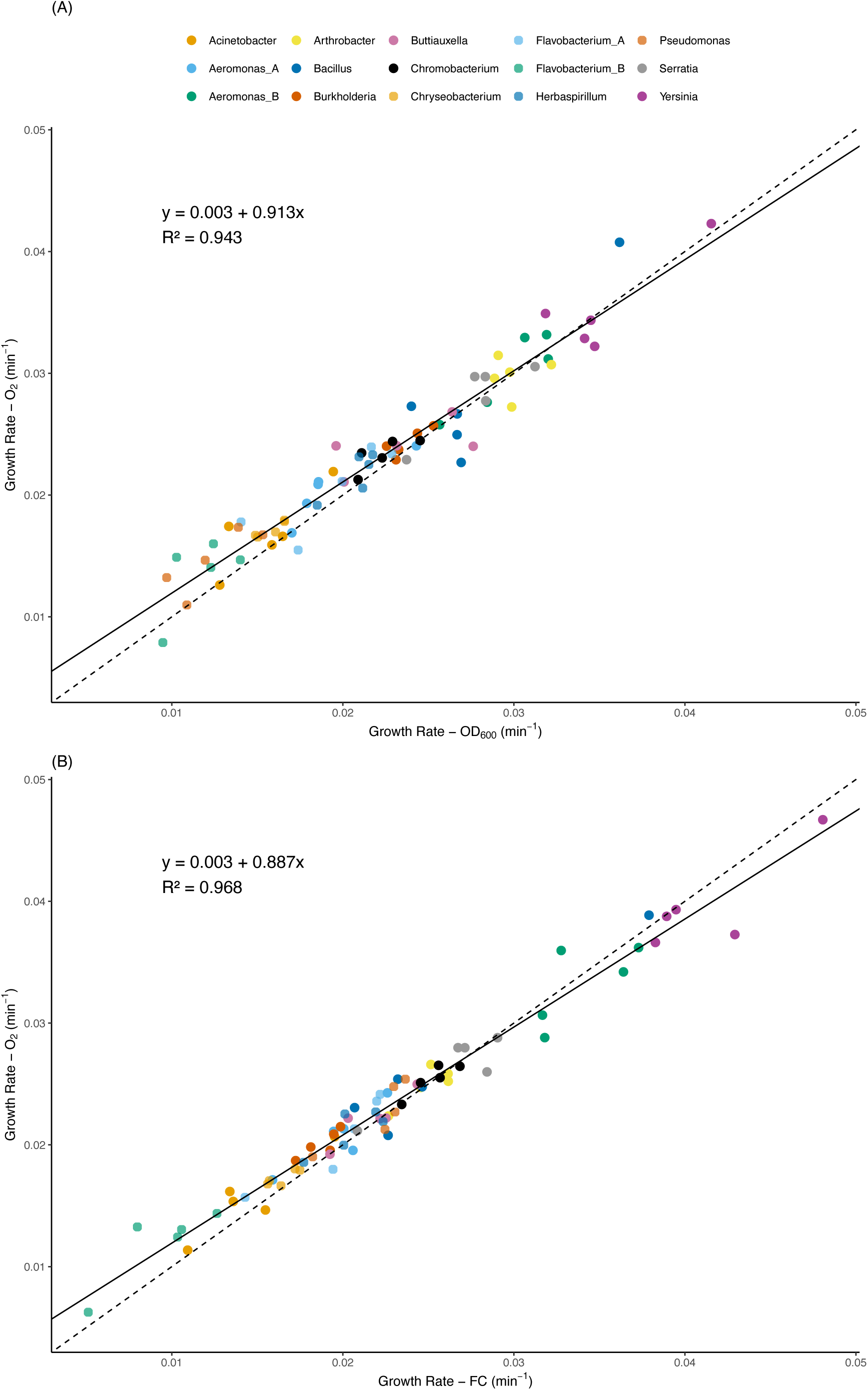
Regression of oxygen-based growth rates against OD₆₀₀ and flow cytometry estimates. Each panel shows replicate-level comparisons between oxygen-derived growth rates (y-axis) and corresponding estimates from OD₆₀₀ (A) or flow cytometry (B) (x-axis). Growth rates were normalised by subtracting taxon-specific random intercepts obtained from linear mixed-effects models. After model fitting, these normalised oxygen-based values were used for plotting. Each point represents a replicate (n = 75; 5 replicates per 15 taxa), coloured by taxon. Black dashed lines indicate the fitted fixed-effect regression lines, while dashed line represent the 1:1 line of equality. Model equations and *R*^2^values are displayed within each panel. Axis limits were kept identical between panels to enable direct comparison of model fit strength and residual variance across methods.

Agreement between oxygen-based and reference methods was evaluated using Bland–Altman analysis (Mansournia et al., 2021) (Fig. 5). For OD₆₀₀-based growth rates (Fig. 5A), oxygen estimates showed a mean bias of 0.001 min⁻¹, with 92.0% of replicate values falling within the limits of agreement (–0.007 to 0.009 min⁻¹). Mixed-effects regression within these limits indicated no evidence of proportional bias (slope = 0.114, *R*^2^= 0.685, *p* = 0.112). Comparisons with flow cytometry yielded similar results (Fig. 5B), with a mean bias of 0.000 min⁻¹ and 97.3% of values within the limits of agreement (–0.004 to 0.005 min⁻¹). The corresponding regression also showed no significant proportional bias (slope = –0.034, *R*^2^ = 0.592, *p* = 0.365).

**Figure 5.**
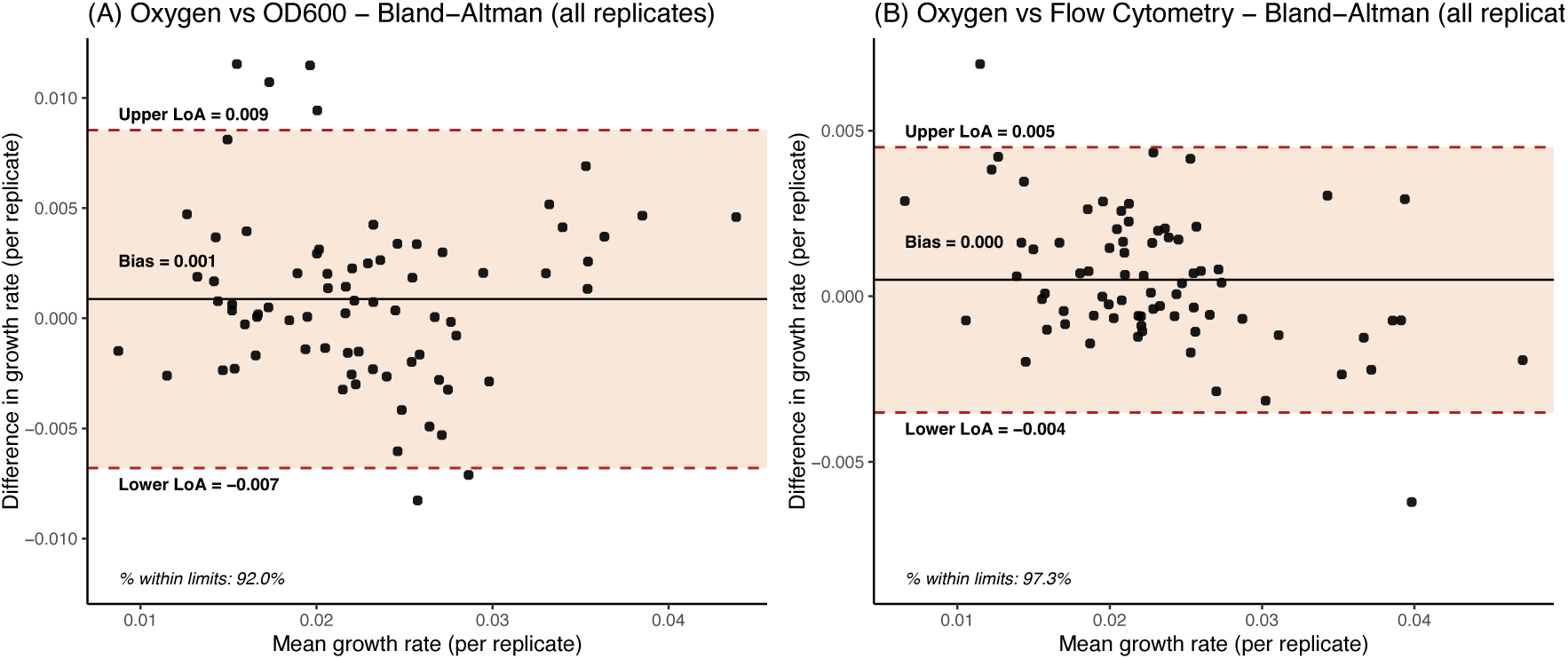
Bland–Altman analysis comparing oxygen-derived growth rates with OD₆₀₀ and flow cytometry estimates. Each panel shows the difference in growth rate estimates between the oxygen-based model and either (A) OD₆₀₀ or (B) flow cytometry (FC) as a function of the mean growth rate for each replicate. Solid horizontal lines indicate the mean bias between methods, while red dashed lines represent the upper and lower limits of agreement (LoA). The percentage of values within these limits is noted in each panel.

### 3.3. Temperature modulation of respiration–growth trade-offs

To investigate how temperature shapes microbial energy allocation, we applied our unified respiration–growth model (eq. 5) to oxygen time series data from *Pseudomonas* sp. exposed to a controlled thermal gradient. Using this framework, we simultaneously inferred growth rate, respiration rate, and carbon use efficiency (CUE), a metric for biosynthetic efficiency (eq. 6) (Fig. 6). Trait–temperature curves were fitted with Sharpe–Schoolfield models for growth and respiration, and the Arrhenius model for CUE, providing activation energies and thermal optima (Padfield et al., 2016; Smith et al., 2021). Growth rates peaked at intermediate temperatures (*T_opt_*= 33.9 °C), consistent with thermal optima for biosynthesis (Fig. 6A), while respiration rates remained elevated at higher temperatures (Fig. 6B; *T_opt_* = 37.5 °C). The activation energies of growth (0.63 eV, 95 % CI 0.57–0.69) and respiration (0.71 eV, 95 % CI 0.59–0.84) overlapped, indicating comparable temperature sensitivities even though respiration peaked at a slightly higher temperature than growth. Carbon-use efficiency (CUE) increased modestly between 20 and ∼30 °C, remained relatively high up to ∼35 °C, and then declined sharply at 40 °C (Fig. 6C), reflecting a shift in energy allocation away from biomass production towards maintenance and cellular repair under high-temperature stress in *Pseudomonas* sp.

**Figure 6.**
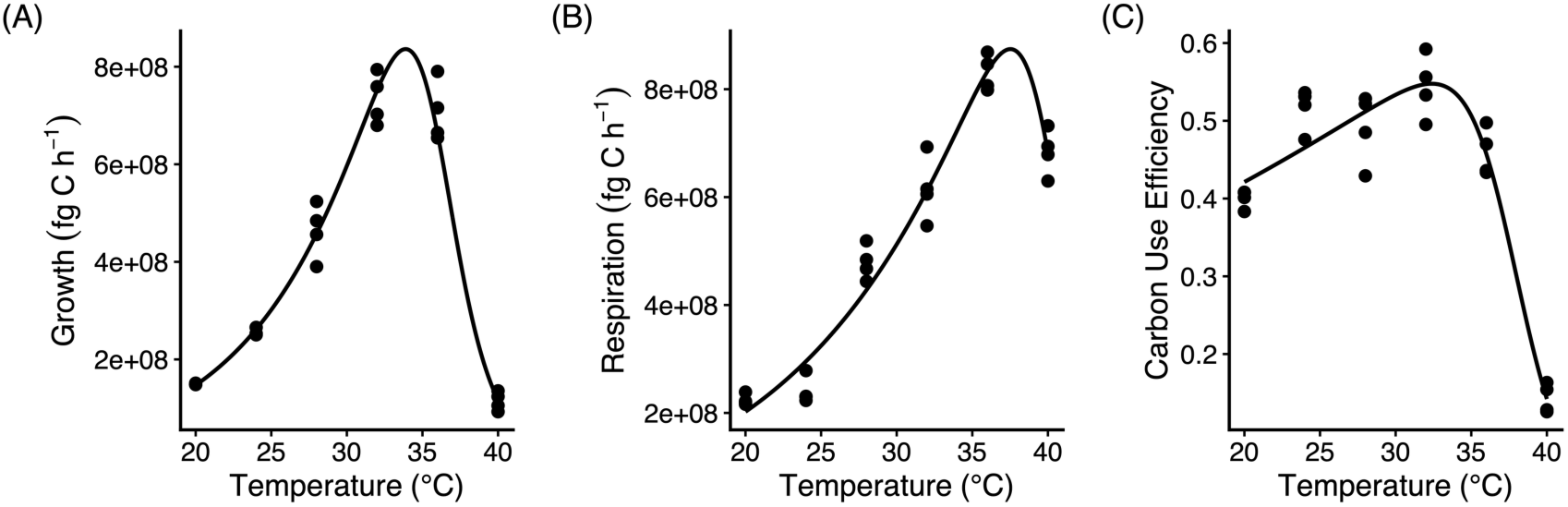
Temperature modulation of respiration–growth trade-offs in Pseudomonas sp. Each panel shows the effect of temperature on microbial energy allocation, inferred from oxygen consumption time series using a unified respiration–growth model. (A) Growth rate, (B) respiration rate, and (C) carbon use efficiency (CUE) are shown across a thermal gradient (20–40 °C). Rates are expressed in femtograms of carbon per hour (fg C min⁻¹), where 1 fg = 10⁻¹⁵ grams. Each point represents an individual replicate. Thermal response curves in (A) and (B) were fitted using the Sharpe–Schoolfield model, which accounts for enzyme kinetics and deactivation at low and high temperatures. In panel (C), an Arrhenius function was used to describe the exponential temperature dependence of CUE over the tested range.

## 4. Discussion

Our results demonstrate that high-resolution oxygen consumption data can be used to estimate bacterial growth and respiration rates simultaneously in diverse bacterial taxa, with strong agreement to independent methods. This represents a major advance because respiration and growth, two of the most fundamental microbial processes, are rarely quantified together from a single, non-invasive assay. By combining growth and respiration within a single analytical pipeline, our framework resolves energy uptake and biomass generation simultaneously and continuously, unlike conventional approaches that rely on parallel assays, often at mismatched temporal resolution (Guo et al., 2022; Lee et al., 2009) and under different physical-chemical conditions. The method captured strain-level variability and produced smooth, monotonic depletion curves with highly consistent trajectories across replicate vials, indicating repeatability and a high signal to noise across diverse taxa.

A key test of the framework is whether our oxygen-based growth estimates agree with independent methods. Growth estimates from the oxygen method showed strong correlation with both flow cytometry and optical density (OD₆₀₀) estimates. The high correlation (*R*^2^> 0.9), low bias and tight limits of agreement in Bland–Altman plots provide strong supporting evidence for the congruence of the oxygen-based growth estimates with traditional. Minor differences between methods were taxon-specific and represented the individual sensitivities of optical, cytometric, and oxygen-based methods (Fig. 3). In particular, slopes slightly below unity (Fig. 4) could reflect modest overestimation of growth by OD- and FC-based methods, as OD may overestimate biomass in taxa that produce extracellular substances (Dalgaard et al., 1994; Koch, 1970), whereas flow cytometry may overestimate biomass by counting all dye-bound, DNA-containing particles, including cellular debris and dead or membrane-compromised cells (Emerson et al., 2017; Zacharias et al., 2015). Our oxygen-based model provides an indirect yet integrated view of metabolic performance and is especially useful for detecting relative differences in microbial traits when biomass production and energy demand diverge. At the same time, agreement among methods does not, by itself, guarantee that the inferred *r* is completely free of bias, because OD₆₀₀-, flow-cytometry–and *O*_2_-based estimates could in principle all be affected by the same deviations from idealised exponential growth (e.g. incipient stationary phase). We therefore interpret the cross-method concordance as strengthening confidence in the *O*_2_-based estimates rather than as absolute proof of their ‘correctness’. Notably, no proportional bias was detected, indicating that concordance between methods is consistent across the full range of growth rates observed.

Independent of methodological considerations, the results highlight distinct ecological patterns in bacterial metabolism. The observed range in growth and respiration strategies suggests bacterial taxa span a spectrum from rapidly growing, energetically demanding phenotypes to slower-growing, low-maintenance organisms (Roller & Schmidt, 2015; Wang et al., 2022; Weissman et al., 2021). This spectrum reflects fundamental life-history trade-offs, directly linking cellular physiology to ecosystem-level processes such as carbon flux partitioning between biomass and respiration. By mechanistically quantifying these traits, our framework provides a pathway to embed microbial physiology into predictive ecosystem models in a way that is both scalable and ecologically realistic.

To demonstrate how the framework can be used to study environmental drivers of microbial metabolism, we applied the method to *Pseudomonas* sp. grown across a 20–40°C temperature range. We found that growth rate was optimal at 33.9 °C, whereas respiration rates continued to rise higher temperatures (see Fig. 6). Consequently, the carbon-use efficiency (CUE), rose gradually up to the optimum temperature and then declined rapidly at high temperature, when respiration rates were high relative to growth rate. These results show how elevated respiration decouples from biosynthesis under heat stress (Apple et al., 2006). By measuring growth and respiration simultaneously from the same oxygen trace, the results reveal how the metabolic costs of maintenance and repair rise more quickly with temperature than biosynthetic capacity, diverting more energy allocation away from biomass synthesis (Kempes et al., 2017).

We emphasise that this temperature-response analysis is best viewed as a proof-of-concept application of the oxygen-based framework rather than a comprehensive description of bacterial thermal physiology. Although the temperature response of CUE aligns with earlier findings, the exact threshold, shape and magnitude of the temperature response will likely differ among taxa with distinct metabolic strategies and thermal niches. These insights from a single model strain connect cellular energetics with functional traits that are important for determining climate-sensitive ecosystem processes and illustrate how the methodology could, in future, be used to quantify microbial responses to warming in high-throughput (Sinsabaugh et al., 2013). This approach could also be applied to investigate other stressors such as osmotic pressure or pH changes, providing a more nuanced view of microbial energy budgeting under changing environmental conditions (Jin & Kirk, 2018; Wood, 2015). More broadly, extending this approach to multiple strains and natural communities will be necessary to characterise the diversity of microbial thermal responses using this novel high-throughput method.

The temperature gradient experiment required expressing growth and respiration in common carbon units, for which we assumed a 1:1 molar O₂:C ratio and a cellular carbon density of 100 fg C µm⁻³. These values are widely used first-order approximations for aerobic respiration of moderately oxidised substrates and for actively growing bacterial cells, but we acknowledge that both O₂:C stoichiometry and cellular carbon content can vary among taxa, substrates, and growth phases. As a result, the absolute carbon fluxes we report for *Pseudomonas* sp. should be interpreted as order-of-magnitude estimates rather than precise stoichiometric measurements. Our main conclusions for the temperature experiment focus on relative responses (how growth, respiration, and CUE change with temperature) rather than on absolute flux magnitudes; moderate shifts in the assumed O₂:C ratio or carbon density would primarily rescale production and respiration and slightly shift CUE values, but are unlikely to overturn the qualitative thermal responses we describe. Future applications could incorporate taxon-specific stoichiometry or formal sensitivity analyses where such information is available.

Several lines of evidence suggest that the patterns we infer are robust to modelling choices and physical confounders. A flexible three-parameter nonlinear function can, in principle, fit many smooth depletion trajectories, so good visual agreement alone does not guarantee that growth and respiration truly remained constant over the full fitting window. Our confidence in the inferred *r* and *R* therefore comes from a broader set of checks rather than curve flexibility per se. Synthetic-data tests show that, under realistic noise levels and sampling schemes, the model reliably recovers both *r* and *R* with very low bias (Supplementary **Table S2**). Consistent with this, the near-perfect agreement between estimates obtained from the standard exponential-decline window and those derived when fits are restricted to high-oxygen data (normalised *O*_2_ ≥ 0.5) indicates that potential low-*O*_2_ limitation has negligible influence on inferred growth and respiration rates (Supplementary **Table S3**). Because initial biomass *N*_0_ only enters when converting the fitted total respiration parameter *R*_tot_ to per-capita respiration, and does not appear in the oxygen time-series model used to estimate *r*, uncertainty in *N*_0_ primarily affects the absolute value of respiration rather than the inferred growth rate (Supplementary **Table S1**). As all oxygen measurements were conducted at constant temperature in headspace-free, gas-tight vials, we expect temperature-driven changes in gas solubility or slow oxygen exchange with the atmosphere to be minimal. The smooth, monotonic depletion curves and highly consistent trajectories across replicate vials are therefore most parsimoniously explained by microbial respiration rather than by physical artefacts such as leakage or pressure-induced oxygen flux. Future studies could refine the current assumptions, for example by using dynamic yield coefficients or explicitly modelling stress-related respiration costs (Schimel et al., 2007), to explore regimes where deviations from exponential growth become more pronounced.

Moreover, combining this oxygen-centric approach with complementary techniques such as CO₂ flux analysis, isotopic tracers, or single-cell methods would provide a multidimensional view of microbial energy use, pushing the field closer to a systems-level understanding of metabolism (Canarini et al., 2020; He et al., 2024; Musat et al., 2016). Overall, we present a scalable system for assessing core microbial metabolic properties using oxygen time series data. By unifying respiration and growth into a single framework, we offer a powerful tool for trait-based microbial ecology, with applications from laboratory phenotyping to ecosystem-scale carbon cycling. Our framework establishes a new benchmark for linking microbial physiology with ecosystem science, opening the way for high-throughput, trait-based mapping of microbial metabolism and providing the physiological bridge needed to predict microbial contributions to global carbon cycling under a rapidly changing climate.

## 5. Conclusions

We present a novel methodological framework that simultaneously estimates bacterial growth and respiration from a single oxygen time series, enabling time-resolved, non-invasive quantification of microbial metabolism. Validated across phylogenetically diverse taxa, the method produced growth estimates consistent with optical density and flow cytometry, demonstrating both congruence with existing methods and broad taxonomic applicability. Critically, by uniting growth and respiration within a single analytical framework, this approach enables direct quantification of their dynamic interaction under controlled conditions, a capability rarely achieved with existing methods.

Application of this framework to a thermal gradient experiment with *Pseudomonas* revealed systematic shifts in respiration-to-growth ratios, demonstrating the method’s capacity to resolve changes in metabolic efficiency along environmentally relevant gradients. These findings illustrate how the approach can detect physiological trade-offs that may be obscured when growth and respiration are measured independently.

This scalable, high-throughput enhances microbial phenotyping by providing integrated metabolic profiles, strengthening trait-based ecology by enabling direct measurement of coupled metabolic processes. As microbial processes become increasingly recognized as central to climate-biosphere feedbacks, such integrative approaches will be essential for embedding mechanistic microbial function into next-generation Earth system models. By quantifying the linkages between growth and respiration across environmental gradients, this framework provides a pathway toward more realistic representations of microbial contributions to global biogeochemical cycles.

## Acknowledgement

We are grateful to the staff of the Living Systems Institute for their help and support.

## Data Availability

The authors confirm that the data supporting the findings of this study are available within the article and its Supplementary Material. Raw data that support the findings of this study are available from the corresponding author, upon request.

## Declaration of generative AI and AI-assisted technologies in the writing process

The authors declare that they have not used any of the generative AI and AI-assisted technologies in this research paper.

## Ethics of research statement

The authors declare that this research paper adheres to high ethical standards of ISME in conducting and reporting the study. All experimental procedures were carried out in accordance with relevant ethical guidelines and regulations.

## Sample CRediT author statement

**Ilgaz Cakin**: Conceptualisation, Methodology, Software, Validation, Investigation, Formal Analysis, Resources, Data curation, Writing - Original Draft

**Rebecca Millington**: Conceptualisation, Methodology, Software, Writing - Reviewing and Editing

**Samraat Pawar**: Conceptualisation, Resources, Supervision, Funding acquisition

**Angus Buckling**: Conceptualisation, Resources, Supervision, Funding acquisition

**Nicholas Smirnoff**: Conceptualisation, Resources, Supervision, Funding acquisition

**Daniel Padfield:** Conceptualisation, Resources, Supervision, Funding acquisition

**John Duffy:** Validation, Resources

**Gabriel Yvon-Durocher**: Conceptualisation, Methodology, Software, Validation, Resources, Writing - Reviewing and Editing, Supervision, Funding acquisition

