## Supplementary Information for "A Novel Method to Simultaneously Estimate Bacterial Respiration and Growth from Oxygen Dynamics"

---

Corresponding author at:
Living Systems Institute, University of Exeter, Stocker Road, Exeter, EX4 4QD, UK

ORCID: 0000-0003-3930-8347
Notes: The authors declare no competing financial interests.

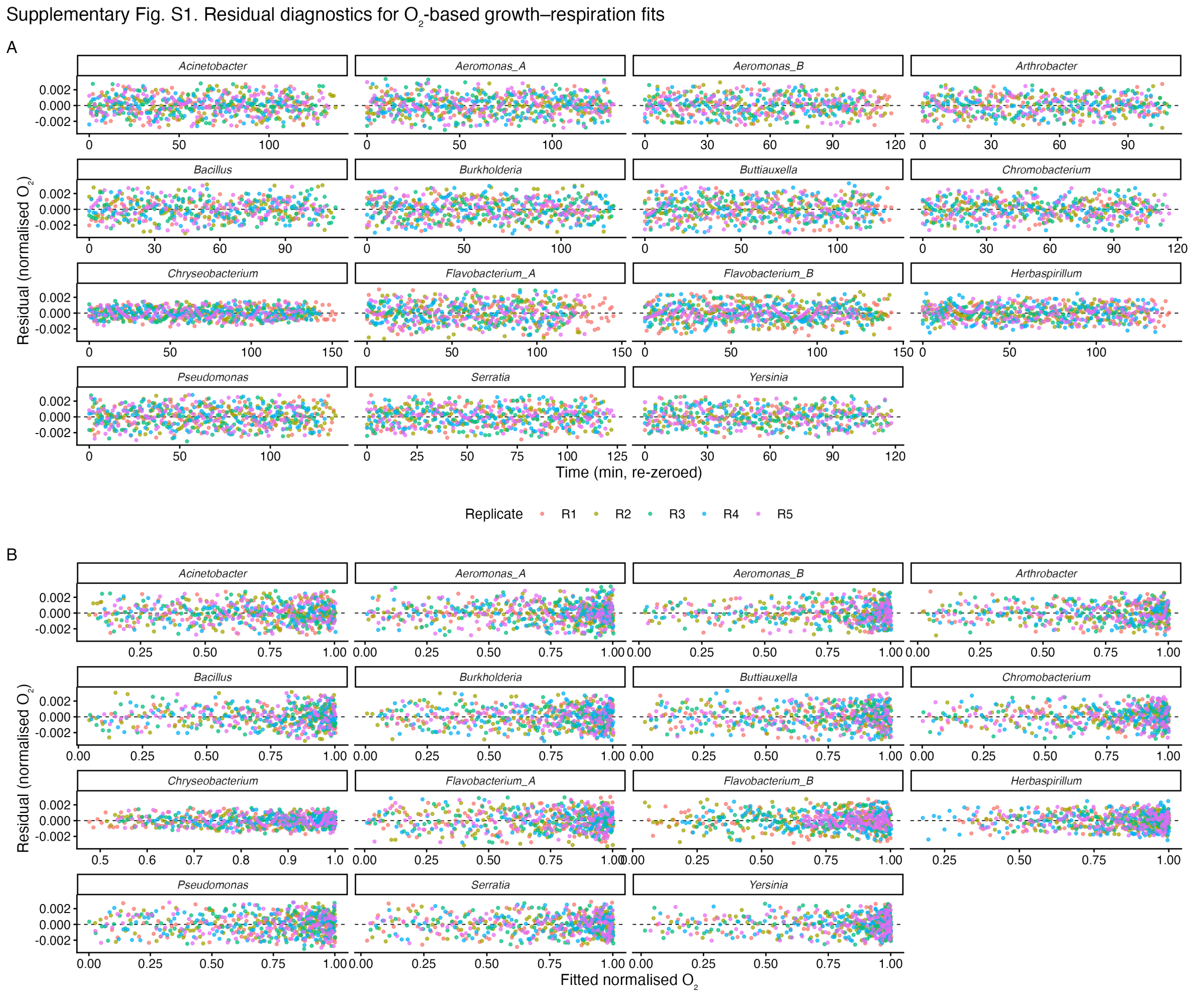


**Supplementary Figure S1.** *Residual diagnostics for $O_2$-based growth–respiration fits. (A) Residuals of normalised $O_2$ as a function of time for each taxon, with points coloured by replicate (R1–R5). (B) Residuals plotted against fitted normalised $O_2$ for the same fits. In both panels residuals are tightly centred around zero with no clear temporal structure or strong heteroscedasticity, indicating that the nonlinear model provides an adequate description of the $O_2$ dynamics across taxa and replicates.*

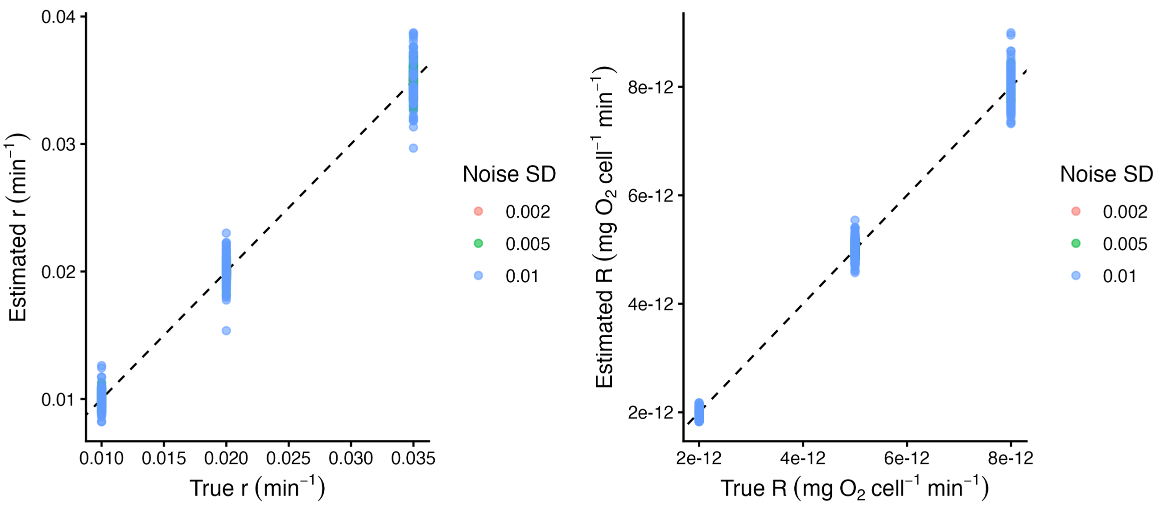


**Supplementary Figure S2.** *Synthetic-data validation of the $O_2$-based growth–respiration model. Left: true vs. estimated growth rate r; right: true vs. estimated per-capita respiration rate R. Each point is a fitted synthetic time series, coloured by the standard deviation of added Gaussian noise (Noise SD); dashed lines show the 1:1 relationship. Across all noise levels, estimates cluster tightly around the 1:1 line, indicating low bias and good parameter recovery for both r and R under realistic experimental noise.*

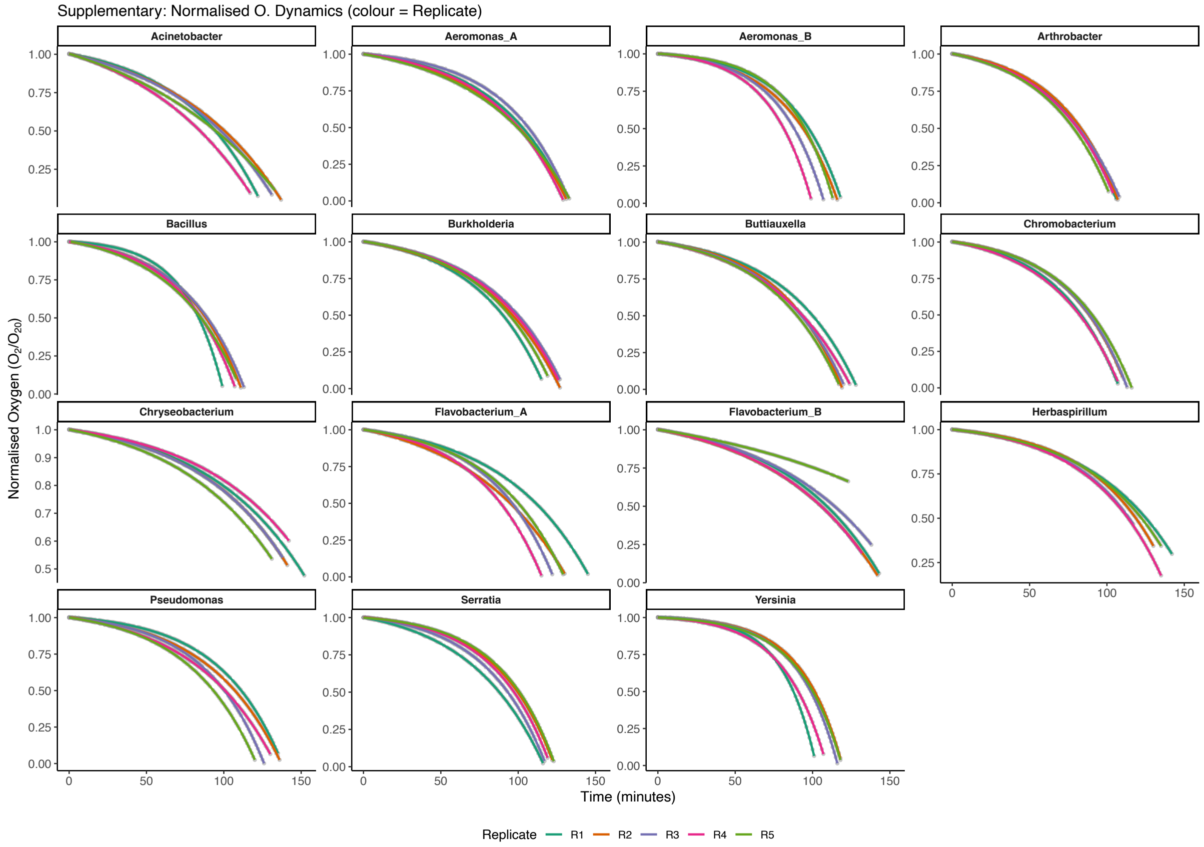

**Supplementary Figure S3.** *Oxygen Dynamics Across All Replicates. Dissolved oxygen concentrations (mg L$^{-1}$) over time (minutes) for each bacterial strain across five replicates (R1–R5). Each panel represents a different strain, with color-coded lines corresponding to individual biological replicates. Smooth lines represent fitted models of oxygen consumption.*

**Supplementary Table S1.** Monte Carlo sensitivity of inferred respiration rates to uncertainty in initial biomass $N_0$. For each taxon, we propagated the empirical variation in $N_0$ across five parallel vials through Monte Carlo resampling (see Methods). The table reports the number of replicates ($n_{\text{reps}}$) and the median, minimum and maximum relative standard deviation (SD/mean) of the resulting Monte Carlo distributions of respiration rate $R$ (columns $R_{\text{rel,sd,median}}$, $R_{\text{rel,sd,min}}$, $R_{\text{rel,sd,max}}$). Relative SDs are generally modest ($<0.35$ for 12 of 15 taxa), with higher propagated uncertainty only for *Bacillus*, *Serratia* and *Flavobacterium* A, where flow-cytometry estimates of $N_0$ were most variable. Growth-rate estimates $r$ were essentially unaffected (median relative SD $\approx 10^{-10}$across taxa; not shown).

| **Taxon** | $n_{\text{reps}}$ | $R_{\text{rel,sd,median}}$ | $R_{\text{rel,sd,min}}$ | $R_{\text{rel,sd,max}}$ |
|---|---|---|---|---|
| *Acinetobacter* | 5 | 0.200 | 0.175 | 0.208 |
| *Aeromonas_A* | 5 | 0.324 | 0.300 | 0.341 |
| *Aeromonas_B* | 5 | 0.287 | 0.265 | 0.301 |
| *Arthrobacter* | 5 | 0.180 | 0.168 | 0.184 |
| *Bacillus* | 5 | 2.654 | 1.676 | 3.299 |
| *Burkholderia* | 5 | 0.116 | 0.113 | 0.119 |
| *Buttiauxella* | 5 | 0.099 | 0.098 | 0.106 |
| *Chromobacterium* | 5 | 0.126 | 0.117 | 0.127 |
| *Chryseobacterium* | 5 | 0.171 | 0.165 | 0.179 |
| *Flavobacterium_A* | 5 | 0.816 | 0.596 | 8.066 |
| *Flavobacterium_B* | 5 | 0.128 | 0.125 | 0.140 |
| *Herbaspirillum* | 5 | 0.214 | 0.204 | 0.228 |
| *Pseudomonas* | 5 | 0.262 | 0.237 | 0.293 |
| *Serratia* | 5 | 1.732 | 0.948 | 6.497 |
| *Yersinia* | 5 | 0.299 | 0.242 | 0.367 |

**Supplementary Table S2.** Synthetic parameter recovery for the normalised oxygen growth–respiration model. For each combination of noise standard deviation, sampling interval and experiment duration, we obtained $n_{\text{fits}}$ successful synthetic fits. Across these fits, we report the bias and RMSE for growth rate $r$ and respiration rate $R$, their relative bias ($r_{\mathrm{rel,bias}}$, $R_{\mathrm{rel,bias}}$), and the mean pseudo-$R^2$ (mean $R^2$).

| noise_sd | dt | duration | $n_{\mathrm{fits}}$ | $r_{\mathrm{bias}}$ | $r_{RMSE}$ | $R_{\mathrm{bias}}$ | $R_{RMSE}$ | $r_{\mathrm{rel,bias}}$ | $R_{\mathrm{rel,bias}}$ | mean $R^2$ |
|---|---|---|---|---|---|---|---|---|---|---|
| 0.002 | 1 | 60 | 450 | 6.5449E-06 | 3.7526E-04 | -1.5373E-15 | 4.7041E-14 | -9.8836E-05 | 7.3552E-06 | 9.9960E-01 |
| 0.002 | 1 | 120 | 450 | 1.8817E-05 | 2.2556E-04 | -4.4278E-15 | 4.2251E-14 | 5.3570E-04 | -5.7188E-04 | 9.9986E-01 |
| 0.002 | 1 | 180 | 450 | 6.6382E-06 | 2.1811E-04 | -1.0993E-15 | 4.0917E-14 | 1.7644E-04 | 3.0360E-05 | 9.9988E-01 |
| 0.002 | 3 | 60 | 450 | -7.9587E-06 | 5.8163E-04 | 1.2187E-15 | 7.2695E-14 | -1.4916E-03 | 2.7937E-04 | 9.9966E-01 |
| 0.002 | 3 | 120 | 450 | 2.0142E-05 | 3.2382E-04 | -2.8458E-15 | 6.0462E-14 | 6.6743E-04 | -7.1734E-04 | 9.9989E-01 |
| 0.002 | 3 | 180 | 450 | 1.7104E-05 | 3.4026E-04 | -2.8748E-15 | 6.2443E-14 | 5.4742E-04 | -6.0953E-04 | 9.9990E-01 |
| 0.002 | 5 | 60 | 450 | 2.3109E-05 | 7.0260E-04 | -3.7178E-15 | 8.1149E-14 | 1.2120E-04 | -5.6114E-04 | 9.9972E-01 |
| 0.002 | 5 | 120 | 450 | -1.5887E-05 | 3.5142E-04 | 1.1292E-15 | 6.9087E-14 | -5.5104E-04 | 1.2072E-03 | 9.9991E-01 |
| 0.002 | 5 | 180 | 450 | 5.4448E-06 | 3.4393E-04 | -1.8073E-15 | 6.7851E-14 | 5.2466E-04 | -5.2179E-05 | 9.9992E-01 |
| 0.005 | 1 | 60 | 450 | 3.2203E-05 | 9.5420E-04 | -1.6320E-15 | 1.1385E-13 | 2.5510E-03 | -8.2889E-04 | 9.9748E-01 |
| 0.005 | 1 | 120 | 450 | 1.3587E-05 | 5.5167E-04 | -3.6286E-15 | 1.0320E-13 | 4.7321E-04 | -2.5448E-04 | 9.9916E-01 |
| 0.005 | 1 | 180 | 450 | 2.8765E-05 | 5.2579E-04 | -6.8108E-15 | 9.9075E-14 | 2.2674E-03 | -4.9943E-04 | 9.9924E-01 |
| 0.005 | 3 | 60 | 450 | -7.0754E-05 | 1.4425E-03 | 6.6218E-15 | 1.7216E-13 | -5.5374E-03 | 3.6635E-03 | 9.9787E-01 |
| 0.005 | 3 | 120 | 450 | -3.5808E-05 | 8.0529E-04 | 1.2110E-14 | 1.5304E-13 | -2.8117E-03 | 1.3195E-03 | 9.9932E-01 |
| 0.005 | 3 | 180 | 450 | -1.8985E-05 | 8.2142E-04 | 3.2595E-15 | 1.5654E-13 | -1.2483E-03 | 1.3970E-03 | 9.9938E-01 |
| 0.005 | 5 | 60 | 450 | 8.1417E-05 | 1.7186E-03 | -1.0020E-14 | 2.1300E-13 | 7.1623E-03 | -1.9109E-03 | 9.9825E-01 |
| 0.005 | 5 | 120 | 450 | 4.0472E-05 | 9.1682E-04 | -4.2089E-15 | 1.8290E-13 | 1.6232E-03 | -5.5252E-04 | 9.9946E-01 |
| 0.005 | 5 | 180 | 450 | 2.5126E-05 | 8.9537E-04 | -4.0475E-15 | 1.8256E-13 | 5.2836E-04 | 6.8679E-04 | 9.9950E-01 |
| 0.01 | 1 | 60 | 450 | -3.5093E-05 | 1.9607E-03 | 5.1486E-15 | 2.4095E-13 | -2.7312E-03 | 3.9817E-03 | 9.9016E-01 |
| 0.01 | 1 | 120 | 450 | -5.7096E-05 | 1.0741E-03 | 1.2566E-14 | 2.0201E-13 | -3.7395E-03 | 3.0265E-03 | 9.9667E-01 |
| 0.01 | 1 | 180 | 450 | -4.0597E-05 | 1.1051E-03 | 9.3503E-15 | 2.0219E-13 | -2.4238E-03 | 2.1317E-03 | 9.9693E-01 |
| 0.01 | 3 | 60 | 450 | -3.1684E-04 | 2.8148E-03 | 6.7042E-14 | 3.6211E-13 | -1.8624E-02 | 1.5216E-02 | 9.9204E-01 |
| 0.01 | 3 | 120 | 450 | -8.9775E-05 | 1.5875E-03 | 1.5261E-14 | 3.0265E-13 | -2.4961E-03 | 2.5461E-03 | 9.9736E-01 |
| 0.01 | 3 | 180 | 450 | -1.3650E-05 | 1.5302E-03 | 1.4784E-14 | 2.9338E-13 | -4.0385E-03 | 1.5927E-03 | 9.9756E-01 |
| 0.01 | 5 | 60 | 450 | 9.7266E-05 | 3.4602E-03 | 4.6912E-15 | 4.1847E-13 | 5.9922E-03 | 3.4401E-03 | 9.9295E-01 |
| 0.01 | 5 | 120 | 450 | 4.1016E-05 | 1.8712E-03 | -1.4575E-15 | 3.6888E-13 | 1.9095E-03 | 8.6902E-04 | 9.9781E-01 |
| 0.01 | 5 | 180 | 450 | -2.3479E-05 | 1.8025E-03 | 1.3429E-15 | 3.6013E-13 | -1.2092E-03 | 2.3208E-03 | 9.9793E-01 |

**Supplementary Table S3.** Sensitivity of fitted parameters to excluding low-oxygen data. For each taxon × replicate, $r_{\mathrm{main}}$ and $R_{\mathrm{main}}$ are the growth rate and per-capita respiration rate estimated from the standard exponential-decline window. $r_{0.5}$ and $R_{0.5}$ are the corresponding estimates obtained when fits are restricted to high-oxygen data (normalised $O_2 \geq 0.5$). The ratio columns give the ratios $r_{0.5}/r_{\text{main}}$ and $R_{0.5}/R_{\text{main}}$, respectively. Values very close to 1 indicate that excluding low-$O_2$ data has only a minor effect on the inferred parameters.

| **Taxon** | **Replicate** | $\boldsymbol{r_{\text{main}}}$ | $\boldsymbol{R_{\text{main}}}$ | $\boldsymbol{r_{0.5}}$ | $\boldsymbol{R_{0.5}}$ | **r ratio** | **R ratio** |
|---|---|---|---|---|---|---|---|
| *Acinetobacter* | R1 | 0.020 | 0.002 | 0.019 | 0.002 | 0.998 | 1.003 |
| *Acinetobacter* | R2 | 0.014 | 0.002 | 0.014 | 0.002 | 1.005 | 0.996 |
| *Acinetobacter* | R3 | 0.015 | 0.002 | 0.015 | 0.002 | 0.997 | 1.002 |
| *Acinetobacter* | R4 | 0.013 | 0.003 | 0.014 | 0.003 | 1.003 | 0.998 |
| *Acinetobacter* | R5 | 0.010 | 0.003 | 0.010 | 0.003 | 0.996 | 1.002 |
| *Aeromonas_A* | R1 | 0.021 | 0.001 | 0.021 | 0.001 | 0.997 | 1.004 |
| *Aeromonas_A* | R2 | 0.020 | 0.002 | 0.020 | 0.002 | 0.997 | 1.004 |
| *Aeromonas_A* | R3 | 0.025 | 0.001 | 0.025 | 0.001 | 1.003 | 0.996 |
| *Aeromonas_A* | R4 | 0.022 | 0.001 | 0.022 | 0.001 | 0.997 | 1.004 |
| *Aeromonas_A* | R5 | 0.018 | 0.002 | 0.017 | 0.002 | 0.995 | 1.004 |
| *Aeromonas_B* | R1 | 0.030 | 0.001 | 0.031 | 0.001 | 1.002 | 0.995 |
| *Aeromonas_B* | R2 | 0.029 | 0.001 | 0.029 | 0.001 | 0.999 | 1.002 |
| *Aeromonas_B* | R3 | 0.034 | 0.001 | 0.034 | 0.001 | 0.998 | 1.004 |
| *Aeromonas_B* | R4 | 0.036 | 0.001 | 0.036 | 0.001 | 0.999 | 1.001 |
| *Aeromonas_B* | R5 | 0.036 | 0.001 | 0.036 | 0.001 | 1.004 | 0.991 |
| *Arthrobacter* | R1 | 0.025 | 0.002 | 0.025 | 0.002 | 1.001 | 0.999 |
| *Arthrobacter* | R2 | 0.026 | 0.002 | 0.026 | 0.002 | 1.000 | 1.000 |
| *Arthrobacter* | R3 | 0.022 | 0.002 | 0.021 | 0.002 | 0.991 | 1.009 |
| *Arthrobacter* | R4 | 0.024 | 0.002 | 0.024 | 0.002 | 0.995 | 1.006 |
| *Arthrobacter* | R5 | 0.024 | 0.002 | 0.024 | 0.002 | 1.006 | 0.993 |
| *Bacillus* | R1 | 0.041 | 0.001 | 0.041 | 0.001 | 0.997 | 1.007 |
| *Bacillus* | R2 | 0.027 | 0.001 | 0.027 | 0.001 | 0.996 | 1.006 |
| *Bacillus* | R3 | 0.025 | 0.002 | 0.025 | 0.002 | 1.000 | 1.001 |
| *Bacillus* | R4 | 0.027 | 0.001 | 0.027 | 0.001 | 0.992 | 1.011 |
| *Bacillus* | R5 | 0.023 | 0.002 | 0.023 | 0.002 | 0.996 | 1.005 |
| *Burkholderia* | R1 | 0.023 | 0.002 | 0.023 | 0.002 | 0.997 | 1.004 |
| *Burkholderia* | R2 | 0.020 | 0.002 | 0.020 | 0.002 | 1.007 | 0.993 |
| *Burkholderia* | R3 | 0.021 | 0.001 | 0.021 | 0.001 | 0.998 | 1.002 |
| *Burkholderia* | R4 | 0.021 | 0.002 | 0.021 | 0.002 | 1.007 | 0.992 |
| *Burkholderia* | R5 | 0.022 | 0.002 | 0.022 | 0.002 | 1.000 | 1.000 |
| *Buttiauxella* | R1 | 0.022 | 0.001 | 0.022 | 0.001 | 1.004 | 0.995 |
| *Buttiauxella* | R2 | 0.024 | 0.001 | 0.025 | 0.001 | 1.004 | 0.994 |
| *Buttiauxella* | R3 | 0.022 | 0.002 | 0.022 | 0.002 | 1.010 | 0.988 |
| *Buttiauxella* | R4 | 0.019 | 0.002 | 0.019 | 0.002 | 0.994 | 1.005 |

| | | | | | | | |
|---|---|---|---|---|---|---|---|
| *Buttiauxella* | R5 | 0.022 | 0.002 | 0.022 | 0.002 | 1.005 | 0.994 |
| *Chromobacterium* | R1 | 0.025 | 0.002 | 0.025 | 0.002 | 0.995 | 1.007 |
| *Chromobacterium* | R2 | 0.026 | 0.001 | 0.026 | 0.001 | 1.001 | 0.999 |
| *Chromobacterium* | R3 | 0.026 | 0.001 | 0.027 | 0.001 | 1.006 | 0.992 |
| *Chromobacterium* | R4 | 0.023 | 0.002 | 0.023 | 0.002 | 1.003 | 0.996 |
| *Chromobacterium* | R5 | 0.025 | 0.001 | 0.025 | 0.001 | 1.004 | 0.994 |
| *Chryseobacterium* | R1 | 0.016 | 0.001 | 0.016 | 0.001 | 1.000 | 1.001 |
| *Chryseobacterium* | R2 | 0.017 | 0.001 | 0.017 | 0.001 | 1.000 | 1.000 |
| *Chryseobacterium* | R3 | 0.017 | 0.001 | 0.017 | 0.001 | 1.000 | 1.000 |
| *Chryseobacterium* | R4 | 0.016 | 0.001 | 0.016 | 0.001 | 1.000 | 1.000 |
| *Chryseobacterium* | R5 | 0.015 | 0.001 | 0.015 | 0.001 | 1.000 | 1.000 |
| *Flavobacterium_A* | R1 | 0.018 | 0.001 | 0.018 | 0.001 | 1.009 | 0.989 |
| *Flavobacterium_A* | R2 | 0.016 | 0.002 | 0.016 | 0.002 | 0.992 | 1.007 |
| *Flavobacterium_A* | R3 | 0.024 | 0.001 | 0.024 | 0.001 | 1.001 | 0.999 |
| *Flavobacterium_A* | R4 | 0.024 | 0.002 | 0.024 | 0.002 | 0.999 | 1.001 |
| *Flavobacterium_A* | R5 | 0.021 | 0.001 | 0.021 | 0.001 | 0.998 | 1.002 |
| *Flavobacterium_B* | R1 | 0.016 | 0.002 | 0.016 | 0.002 | 1.007 | 0.994 |
| *Flavobacterium_B* | R2 | 0.015 | 0.002 | 0.015 | 0.002 | 1.002 | 0.998 |
| *Flavobacterium_B* | R3 | 0.014 | 0.002 | 0.014 | 0.002 | 0.998 | 1.002 |
| *Flavobacterium_B* | R4 | 0.015 | 0.002 | 0.015 | 0.002 | 1.001 | 1.000 |
| *Flavobacterium_B* | R5 | 0.008 | 0.002 | 0.008 | 0.002 | 1.000 | 1.000 |
| *Herbaspirillum* | R1 | 0.018 | 0.001 | 0.018 | 0.001 | 0.999 | 1.001 |
| *Herbaspirillum* | R2 | 0.023 | 0.001 | 0.023 | 0.001 | 1.000 | 1.000 |
| *Herbaspirillum* | R3 | 0.022 | 0.001 | 0.022 | 0.001 | 0.999 | 1.001 |
| *Herbaspirillum* | R4 | 0.022 | 0.001 | 0.022 | 0.001 | 0.995 | 1.008 |
| *Herbaspirillum* | R5 | 0.020 | 0.001 | 0.020 | 0.001 | 1.001 | 0.999 |
| *Pseudomonas* | R1 | 0.025 | 0.001 | 0.025 | 0.001 | 1.003 | 0.995 |
| *Pseudomonas* | R2 | 0.021 | 0.001 | 0.021 | 0.001 | 0.997 | 1.003 |
| *Pseudomonas* | R3 | 0.025 | 0.001 | 0.025 | 0.001 | 0.996 | 1.007 |
| *Pseudomonas* | R4 | 0.019 | 0.002 | 0.019 | 0.002 | 1.001 | 0.999 |
| *Pseudomonas* | R5 | 0.023 | 0.002 | 0.023 | 0.002 | 1.006 | 0.993 |
| *Serratia* | R1 | 0.021 | 0.002 | 0.021 | 0.002 | 1.004 | 0.996 |
| *Serratia* | R2 | 0.028 | 0.001 | 0.027 | 0.001 | 0.998 | 1.004 |
| *Serratia* | R3 | 0.026 | 0.001 | 0.026 | 0.001 | 1.002 | 0.998 |
| *Serratia* | R4 | 0.028 | 0.001 | 0.028 | 0.001 | 0.998 | 1.002 |
| *Serratia* | R5 | 0.028 | 0.001 | 0.028 | 0.001 | 1.001 | 0.998 |
| *Yersinia* | R1 | 0.046 | 0.000 | 0.046 | 0.000 | 1.000 | 1.000 |
| *Yersinia* | R2 | 0.038 | 0.000 | 0.038 | 0.000 | 1.002 | 0.996 |
| *Yersinia* | R3 | 0.039 | 0.000 | 0.039 | 0.000 | 0.996 | 1.010 |

| | | | | | | | |
|---|---|---|---|---|---|---|---|
| *Yersinia* | R4 | 0.037 | 0.001 | 0.037 | 0.001 | 0.999 | 1.001 |
| *Yersinia* | R5 | 0.036 | 0.000 | 0.036 | 0.001 | 0.999 | 1.003 |

**Supplementary Note S1**. Practical guidance for implementing the oxygen-based respiration–growth model

Through repeated experiments and model fitting, we identified several common pitfalls and developed a set of practical solutions to ensure reproducibility of oxygen-based respiration–growth measurements. We summarise these here to provide a step-by-step guide.

**Culture preparation.** The most frequent issue is that cultures are either too dense or still in lag phase at the start of the assay. If biomass is too high, oxygen becomes exhausted before the exponential phase can be captured. Conversely, if cells remain in lag phase, the growth signal is weak and parameter estimates are unreliable. To avoid both, we recommend pre-growing cultures for approximately two hours at $OD_{600}$ = 0.05 to ensure they exit lag phase, and then diluting to $OD_{600} \approx 0.0005$ in PreSens vials. This typically provides at least 60 minutes of measurable, exponential oxygen depletion, sufficient for robust fitting.

**Loading vials.** Oxygen re-equilibration and sensor drift often arise from poor vial handling. Bubbles and headspace permit re-oxygenation, while leaky caps allow slow drift. To prevent this, vials should be overfilled to eliminate headspace and capped tightly. After capping, swirl gently to dislodge bubbles before placing vials onto the reader.

**Reader setup.** Sensor spots are sensitive to environmental conditions such as drift, temperature fluctuations, and ambient light. To minimise these effects, use factory-calibrated sensor spots or perform a two-point calibration before each experiment. Plates should then be allowed to stabilise for 25–30 minutes at the assay temperature (e.g., 30 °C) before logging begins.

**Measuring initial biomass ($N_0$).** Accurate estimation of $N_0$ is critical, as errors in starting density bias both the growth rate $(r)$ and respiration rate $(R)$. However, sampling at $t_0$ is easily disrupted by opening the incubator door. A practical workaround is to prepare parallel vials in a separate incubator for $N_0$ measurement. Biomass can be quantified by flow cytometry or OD, and these values should be supplied directly into the model as $N_0$.

**Model fitting.** Non-convergence occasionally occurs, especially if starting values are poorly chosen. If this happens, manually adjusting starting values (e.g., for $r$ or $R$) generally resolves the issue. In addition, we recommend constraining parameters to broad, physiologically motivated intervals that act as guardrails to exclude clearly non-physical solutions without tightly restricting the fit. In our analyses, the specific growth rate was bounded by $1 \times 10^{-4} \leq r \leq 0.1\ \text{min}^{-1}$, corresponding to doubling times from ~7 h to ~7 min. The per-cell respiration rate was bounded by $1 \times 10^{-12} \leq R \leq 1 \times 10^{-6}$mg $O_2$ cell$^{-1}$ min$^{-1}$, a range several orders of magnitude wider than the values typically recovered from our cultures. For normalised oxygen series, the initial oxygen term was constrained to $0.8 \leq O_{2,0} \leq 1.2$, allowing ±20% deviation around the nominal starting value after normalisation. These bounds help keep the optimisation within realistic regions of parameter space while leaving the inferred parameters largely driven by the data.

**Fit diagnostics.** Visual inspection alone can be misleading: some curves may appear to fit but yield poor statistical support, or patterned residuals. We recommend applying a pseudo-$R^2$

threshold of ≥ 0.995, based on observed distributions (median = 0.99997 in our data). Fits that exhibit strong residual patterns, particularly at early or late time points, should be discarded.

**Replication.** Finally, insufficient replication reduces confidence and makes it difficult to detect sensor drift or outliers. We advise using at least four biological replicates per treatment, alongside blanks that allow monitoring of baseline drift in sensor readings.
